# The breast pre-cancer atlas illustrates the molecular and micro-environmental diversity of ductal carcinoma in situ

**DOI:** 10.1101/2021.05.11.443641

**Authors:** Daniela Nachmanson, Adam Officer, Hidetoshi Mori, Jonathan Gordon, Mark F. Evans, Joseph Steward, Huazhen Yao, Thomas O’Keefe, Farnaz Hasteh, Gary S. Stein, Kristen Jepsen, Donald L. Weaver, Gillian L. Hirst, Brian L. Sprague, Laura J. Esserman, Alexander D. Borowsky, Janet L. Stein, Olivier Harismendy

**Affiliations:** Bioinformatics and Systems Biology Graduate Program, University of California San Diego; Division of Biomedical Informatics, Department of Medicine, University of California San Diego; Department of Pathology and Laboratory Medicine, Center for Immunology and Infectious Diseases, School of Medicine, University of California Davis; University of Vermont Cancer Center; Department of Biochemistry, University of Vermont; Department of Pathology and Laboratory Medicine, University of Vermont; Moores Cancer Center, University of California San Diego; Institute for Genomic Medicine, University of California San Diego; Department of Surgery, University of California San Diego; Department of Pathology, University of California San Diego; Helen Diller Family Comprehensive Cancer Center, University of California San Francisco; Department of Surgery, University of Vermont

## Abstract

Micro-environmental and molecular factors mediating the progression of Breast Ductal Carcinoma In Situ (DCIS) are not well understood, impeding the development of prevention strategies and the safe testing of treatment de-escalation. We addressed methodological barriers and characterized the mutational, transcriptional, histological and microenvironmental landscape across 85 multiple micro-dissected regions from 39 cases. Most somatic alterations, including whole genome duplications, were clonal, but genetic divergence increased with physical distance. Phenotypic and subtype heterogeneity frequently associated with underlying genetic heterogeneity and regions with low-risk features preceded those with high-risk features according to the inferred phylogeny. B- and T-lymphocytes spatial analysis identified 3 immune states, including an epithelial excluded state located preferentially at DCIS regions, and characterized by histological and molecular features of immune escape, independently from molecular subtypes. Such breast pre-cancer atlas with uniquely integrated observations will help scope future expansion studies and build finer models of outcomes and progression risk.

## Introduction

Increasing adoption of breast cancer screening and advances in imaging capabilities have improved our ability to identify breast ductal carcinoma in situ (DCIS). Rarely diagnosed 40 years ago, DCIS now comprises nearly 20% of all breast cancer-related diagnoses ^1, 2^. Unfortunately, this progress has not resulted in decreased breast cancer mortality. Standard treatment, involving surgical excision often complemented with radiation therapy (in the setting of breast conserving surgery) and endocrine recurrence risk reduction (particularly with ER+ DCIS), therefore constitutes overtreatment, and not without treatment-related consequences for many ^2, 3^. DCIS progression is particularly difficult to study longitudinally due to the current standard of surgical excision of the lesion and the infrequent progression and/or occurrence of new primary lesions over a long timespan(5-10% after 10 years) ^4^. Clinicopathological risk factors such as large size, dense breast, younger age, high pathological grade, presence of comedo-necrosis or Her2 positivity have been associated with increased risk of recurrence, but the resulting predictive models, or those relying on gene expression signatures, are currently insufficient to safely distinguish patients to watch from patients to treat ^5^.

Contrary to common conceptual models, there is little evidence for the sequential accumulation of somatic alterations during progression from in situ to invasive disease, but rather all invasive breast cancer (IBC) intrinsic subtypes and known driver mutations have been identified in DCIS, albeit at variable prevalence ^6–12^. Moreover, both single-cell and bulk studies have shown similar clonal make-up of synchronous invasive and in situ lesions, convoluting the idea that clonal selection drives invasion ^11, 13^. The role of the immune environment has also been investigated, highlighting the higher lymphocyte infiltration in Her2+ or Triple Negative DCIS, or specific immunological make-up of samples at higher risk of progression ^14–16^. Similarly, the role of the basal layer, fibroblasts, adipocytes, other stromal cells or overall extracellular matrix has identified features that are different between DCIS and IBC, likely mediated by chemokine signalling and can be associated with known progression risk factors ^17–20^. Their active participation in the malignant transformation of the breast epithelium remains to be established as similar mechanisms are typically involved in normal development, activity and aging of the mammary gland ^21^. ^12^

Progress in our understanding of the processes mediating DCIS onset and progression has been considerably hindered by technical and logistical limitations. Indeed, pure DCIS lesions are commonly small in size, formalin and paraffin embedded (which damages nucleic acids) and can display significant histological heterogeneity ^22^. As a consequence, comprehensive molecular and cellular assays and their integrated analysis have seldom been performed in pure DCIS cohorts. Capturing evidence of phenotypic, genetic and cellular heterogeneity, and how they relate to each other is necessary to develop a better spatial, temporal and functional understanding of the mechanisms at play. Recent advances in genome-wide assays, becoming compatible with ever more challenging samples ^23–26^, have improved our ability to connect histological and molecular observations and enabled such application even to individual microbiopsies from a histological slide of pure DCIS.

Here we describe the combined, parallel histological, molecular and immunological profiling of pre-malignant lesions from 39 patients diagnosed with DCIS, including multiple epithelial micro-biopsies within a subset of samples. The dissection of specific epithelial lesions provided a detailed assessment of the association of their histological architecture with intrinsic subtypes, mutational landscape, driver mutations and immunological states. Multi-region profiling resulted in the inference of clonal relationships, illustrating how genotypes related to phenotypes within a specimen. To our knowledge, our report is the first multi-modal and sub-histological profiling of a cohort of pure DCIS, illustrating spatial heterogeneity and placing diverse states of immune-activity observed in their specific molecular and histological context.

## Results

### Histological and molecular characterization

We collected a total of 43 specimens (referred to as samples) from 39 patients diagnosed with pure DCIS, including three samples from subsequent DCIS diagnosed between 14 and 70 months after the index DCIS (Figure 1a-b, Table S1). Sixty-nine percent (29/42) of the samples were positive for estrogen receptor (ER) expression and 40% (16/40) had *ERBB2* gene overexpression or amplification (Figure S1a). Each sample was further annotated for grade and histological architecture and the annotations were used to identify regions of interest, guide the micro-biopsies of the epithelial areas and the immuno-histological analysis. On the basis of their studied regions, the cohort consisted of 32 high or intermediate grade DCIS (HG-DCIS), 9 low-grade DCIS (LG-DCIS) and 2 low-grade atypical ductal hyperplasia (ADH). The DCIS regions could be further annotated according to their dominant histological architecture (17 cribriform, 19 solid, 3 mixed, 2 micropapillary) and the presence of necrosis (10 comedo-necrosis, 17 other). LG-DCIS were more frequently of cribriform architecture (8/9), while HG-DCIS were frequently necrotic (25/32). The relative area of adipose tissue in each sample varied between 4 and 91 percent as estimated by segmental classification of the whole slide digital image (Figure 1c, Methods). The lower adipose fraction was associated with higher mammographic breast density (p=0.0067) suggesting the sample histology was representative of the whole breast texture. Interestingly, solid DCIS were associated with a higher adipose fraction (median 69% vs 40%, p=0.008), suggesting a contribution of the breast microenvironment to the growth architecture. Overall, the cohort represents a diverse set of pure in situ lesions identified in absence of any detectable invasive component. The studied samples are enriched for DCIS lesions and specifically annotated for their histological architecture. Each sample was profiled using multiple assays, performed on sequential histological sections (4-7 µm) used for whole transcriptome, whole-exome and spatial immune profiling. Whenever possible, the investigated regions were matched across assays to preserve the spatial information in the analysis and limit the variation due to spatial heterogeneity. Spatial heterogeneity was further addressed in 21 samples for which multiple sub-regions were profiled independently.

**Figure 1:**
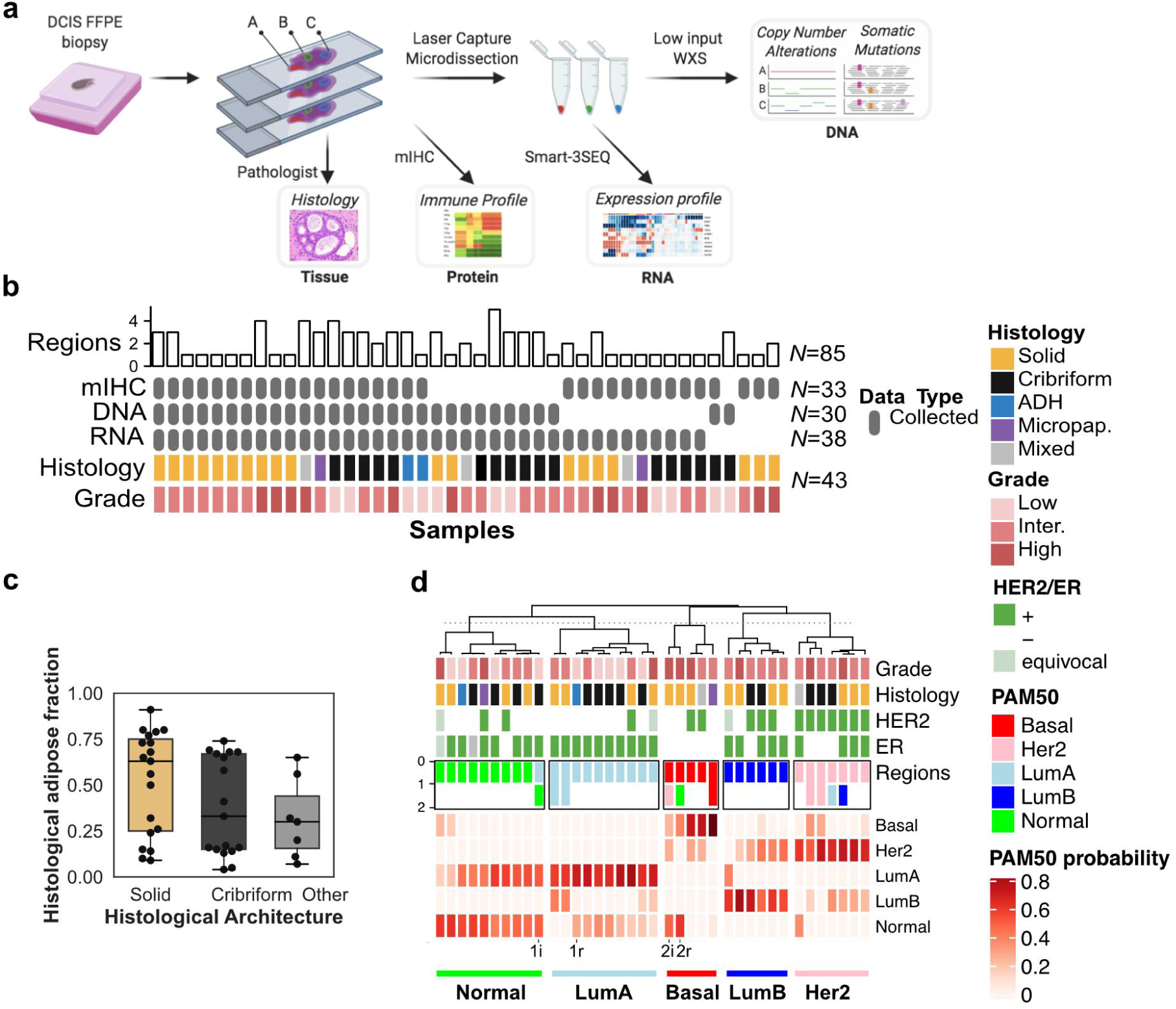
Study design and cohort overview. **(a)** Archival sample processing and analysis workflow including histology (H&E and mIHC) and microbiopsy-derived whole exome (WXS) and whole transcriptome (Smart-3SEQ) profiling. **(b)** Study cohort overview including histological characteristics (colored rows), data type (grey rows) and number of histological regions (bar chart) investigated. **(c)** Estimate of the fraction of adipose area in H&E images in epithelium of the mIHC images according to each histological architecture. **(d)** Sample classification according to the probabilities of each PAM50 expression subtype. For 10 eligible samples, the intrinsic subtype of a spatially distinct region is indicated. Two patients with recurrence (r) and index (i) samples are indicated at the bottom.

The expression of genes was measured using high-throughput sequencing of RNA-seq libraries directly prepared from the micro-biopsied regions ^24^ (Table S2). The samples were classified according to the PAM50 intrinsic subtypes used for invasive breast cancer (IBC), which identified Basal (N=5), Luminal A (N=10), Luminal B (N=6), Her2-like (N=7) and Normal-like (N=10) samples (Figure 1d). Consistent with IBC classification, Luminal A and B were enriched for samples from ER+ cases, while Her2-like were enriched for Her2+ cases. Similarly, Luminal B and Her2-like were enriched in HG-DCIS, while Luminal A was almost exclusively composed of cribriform LG-DCIS. Luminal A and Normal-like represented closely related classes and together comprised the majority of the samples (20/38), which is not unexpected given the higher fraction of low-grade and pure in-situ lesions in the cohort, in contrast with IBC and previous DCIS expression profiling studies ^27, 28^. The PAM50 subtype of two independent sub-regions with matching histology and grade was determined in 10 samples, and observed to be discordant in 5 samples (Table S2B), which was associated with larger distances between the regions (Mann Whitney, p=0.005, Figure S1b). Interestingly, matched index and recurrent samples from two patients had at least one region with concordant subtype. Across all samples the distribution of probabilities for each PAM50 subtype likely captures such heterogeneity. Normal-like were truly a mix of Normal and Luminal A, while Her2-like tended to have two main subsets: Her2/Basal and Her2/Luminal B. This suggests that subtypes inferred from bulk analysis, even after epithelial micro-dissection, are frequently the result of a variable mixture of pure subtypes.

### Subtype differences in mutational landscape

To determine whether any of the histological or molecular subtypes described above were associated with specific genetic alterations, we characterized their mutational landscape. Whole exome sequencing was carried out on micro-biopsies from 30 samples using a procedure specifically optimized for low amount of damaged DNA ^23^. Mutations and copy number alterations (CNA) were identified in 27 and 30 samples, respectively (Table S3). The median copy number burden - or fraction of the genome involved in CNA – was 0.14 and was 2.5 fold higher in HG-DCIS (Mann Whitney, p=0.017, Figure 2a-b). Whole genome doubling (WGD) events were detected in 3/8 eligible samples, all of which were low or intermediate grade cribriform DCIS consistent with its early timing in breast carcinogenesis ^29^ (Table S3). Consistent with previous studies, loss of 16q (13/30) and 17p (12/30) or gain of 1q (12/30) were among the most frequent chromosomal alterations and, while many events were more frequent in HG-DCIS (Figure 2c, Table S4), these hallmarks were also observed in low-grade or benign lesions, including ADH: (1q gain: 1/9, 16q loss: 7/9, 17p loss: 3/9).

**Figure 2.**
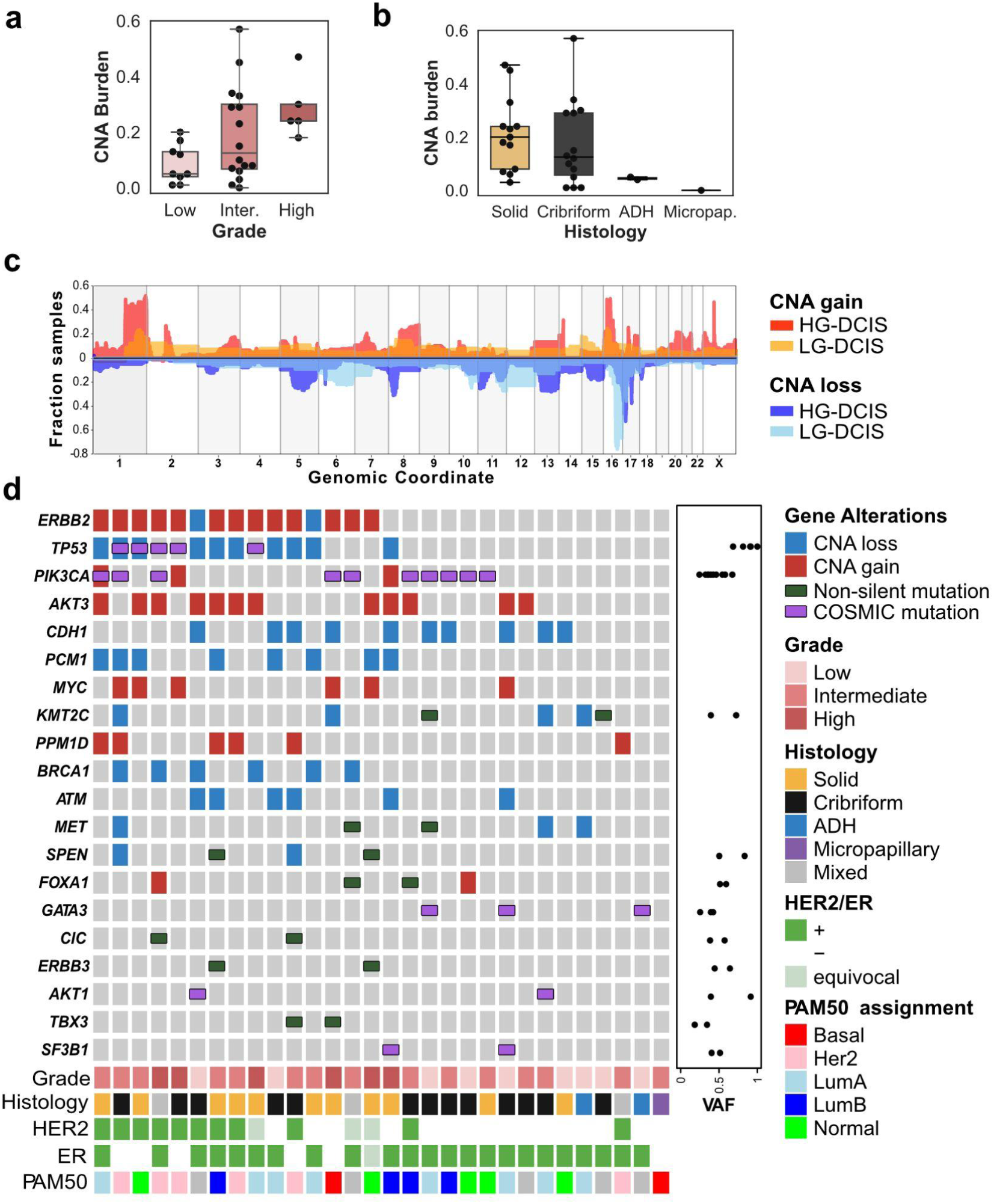
Pure DCIS genomic landscape. **(a-b)** CNA burden (fraction of base pairs involved in copy number gain or loss) as a function of grade (a) and (b) histological architecture. **(c)** Smoothed frequency (y-axis) of CNA gains (top) and losses (bottom) smoothed along the genome (x-axis) for HG-DCIS (N=22 - dark colors) and LG-DCIS (N=5 light colors). **(d)** Oncoprint diagram displaying the mutational status of driver genes commonly altered in breast cancer. Genes were included if they were mutated in at least 2 patients or located in a CNA segment present in at least 6 patients, and ordered by frequency of alteration. The variant allele fraction (VAF) of mutations (right panel) and histological characteristics (bottom panel) are indicated.

We identified between 74 and 207 coding mutations per sample. The mutational burden was higher in HG-DCIS (Mann Whitney, p=0.003) and Her2-like subtypes (Mann Whitney, p=0.025), recognizing that these categories are overlapping. The HG-DCIS burden (4.4 mut/Mb) was higher than previous reports, possibly due to residual germline variants in our study ^11, 13^. We identified aging-associated mutational signatures (SBS1 and SBS5) in all samples eligible for analysis (N=13), APOBEC signature (SBS2 and SBS13) on 1 intermediate grade solid DCIS and mismatch repair signature (SBS15 or SBS21) in 3 DCIS of variable grade and architecture (Table S5). The APOBEC signature is therefore more rare in DCIS than IBC (∼8% vs >75%), but can be present in premalignant lesions. Interestingly, this sample also displayed clustered mutations (N=3 within 1,416 bp) in chromosome 17q (Figure S2), an APOBEC-driven kataegis site frequently seen in IBC ^30^. The most recurrently mutated genes were *PIK3CA* (44%), *TP53* (31%), and *GATA3* (20%), and were all affected by known somatic mutations in breast cancer at similar rates to previous studies of pure DCIS ^6, 7, 9–11^ (Figure 2d & Table 1). *TP53* mutations were only found in HG-DCIS and associated with high CNA burden (Mann-Whitney, p=0.018), while *GATA3* mutations were only found in cribriform or ADH histologies and associated with LG-DCIS (Fisher Exact, p=0.005). Interestingly, *GATA3* mutations were identified in larger lesions (Mann Whitney, p=0.038), consistent with a similar observation in invasive cancer and the larger tumor size of *GATA3* mutated xenograft models ^31, 32^. Another 9 selected genes known to be mutated in IBC were recurrently affected by 18 mutations predicted to be deleterious, 4 of which are known somatic mutations ^7^. The result suggests that oncogenic driver mutations are already present at the premalignant stage, including in LG-DCIS (e.g. *SF3B1* c.2098A>G) or ADH (*GATA3* c.925-3_925-2del). This is consistent with previous reports and reports of field effect mutations in normal ducts or benign lesions ^33, 34^, though the contribution of these mutations to the lesion progression remains to be determined.

**Table 1.**
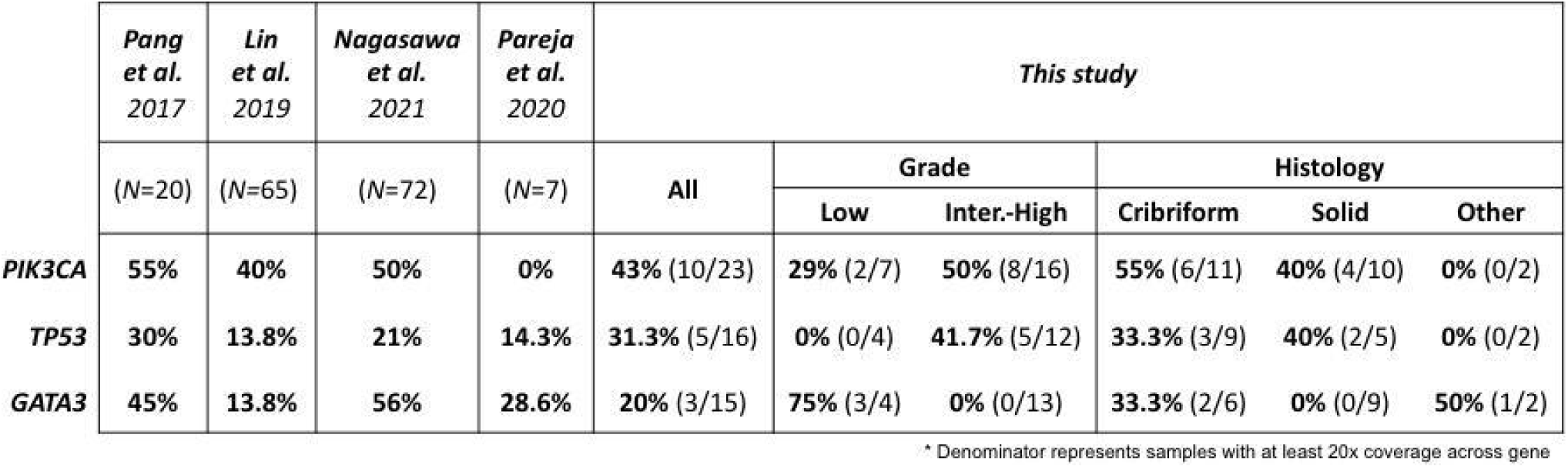
Frequency of *PIK3CA*, *TP53* and *GATA3* driver mutations in previously reported (Pang et al., Lin et al., Nagasawa et al., Pareja et al.) and current pure DCIS cohort.

### Genetic heterogeneity and clonal diversity

The histologic assessment and expression profiling have revealed variable levels of phenotypic heterogeneity across the samples. In order to determine whether such heterogeneity is present at the genetic level, we measured genetic heterogeneity in two distinct ways: 1) divergence, which measures the genetic distance between regions of a sample and, 2) clonal relationships, which uses phylogenetic tree construction to establish evolutionary order to genetic alterations (Table S3). We measured divergence by computing a CNA-based score on 19 pairs of histologically matching regions in 11 samples (Table S3, Methods). With no pairs completely independent (max score=0.16), the spatial distance separating the dissected regions was correlated with the extent of their genetic divergence (R^2^=0.65, p=0.00017, Figure S3), suggesting that, locally, genetic diversity results from wider range proliferation. Divergence was the lowest in 1 ADH, but was not associated with grade, Her2/ER status, or adipose fraction suggesting that local genetic heterogeneity is not associated with progression risk factors.

More precise clonal relationships between regions were evaluated using phylogenetic analysis in 12 samples, comparing CNA, and mutations when available (Figure 3, Figure S4, Methods). While the majority (88.4%) of CNA were shared across all regions of a sample, 11.6% were private to some regions, as observed in 7/12 samples. Multiple samples (3/12) contained mutations in putative cancer driver genes that were private to one region only. These included known and likely pathogenic mutations in *ATR, PIK3CA, MET, KDM5C,* suggesting that not all driver mutations are acquired early. Interestingly, the three samples with the most private CNA displayed discordant histological architecture or discordant PAM50 subtypes between regions, suggesting that within a sample, genetic and phenotypic differences are linked. Furthermore, in 4/5 samples containing regions with discordant histology and 3/4 with discordant PAM50 subtypes, benign regions or regions with Normal or LumA subtypes appeared earlier than regions with Her2 or Basal subtypes or regions with necrosis. This result suggests that in these samples the lesion progressed with time and acquired histological and molecular features previously associated with increased risk of progression.

**Figure 3.**
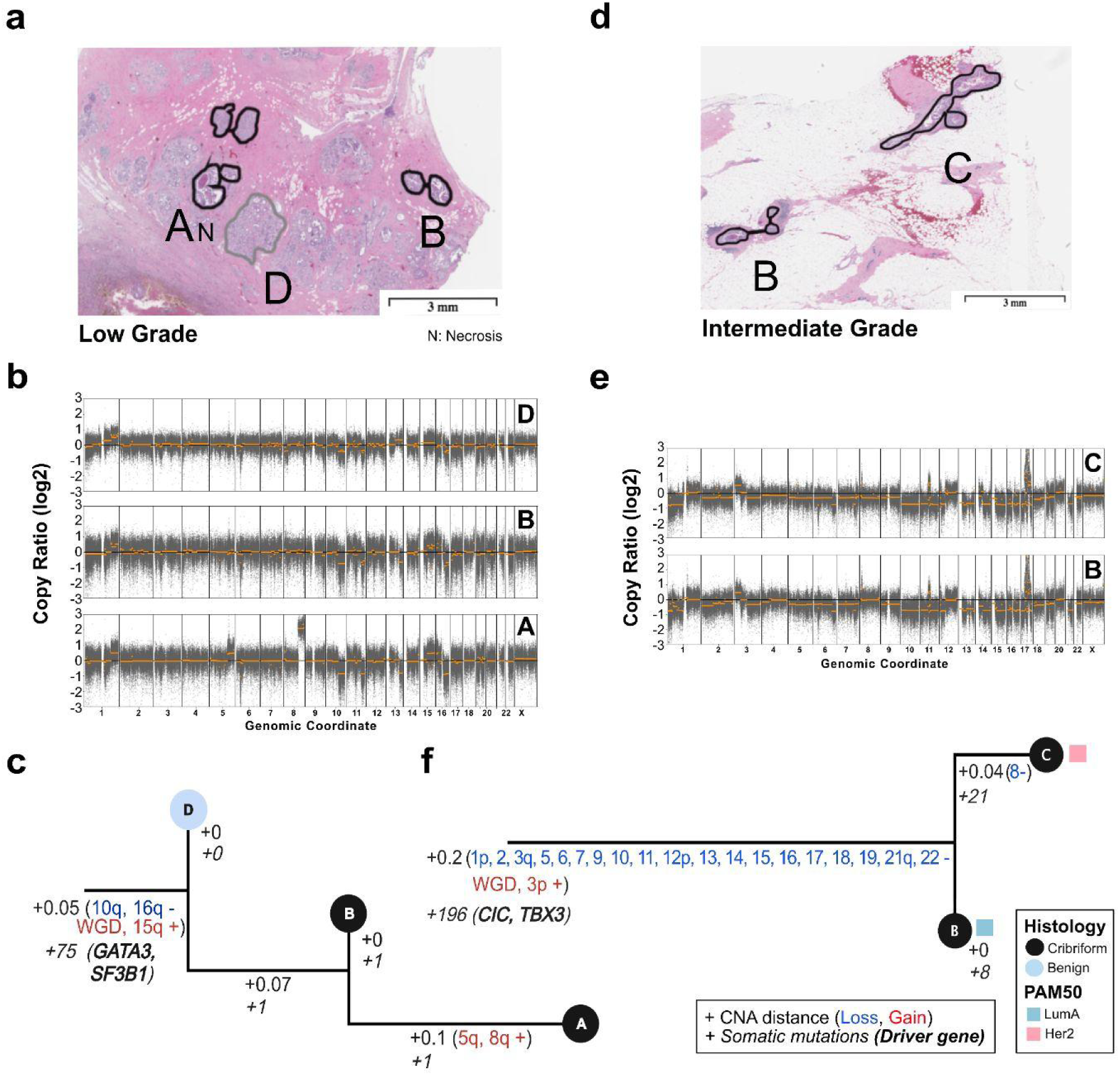
Clonal relationships of multi-region DCIS. Multi-region phylogenetic reconstruction using both CNA and somatic mutations for MCL76_061_16200 **(a-c)** and MCL76_077_15300 **(d-f)**. For each case, the spatial annotation of the microdissected regions on the H&E images (a,d), corresponding copy number profiles (b,e) and phylogenetic trees (c,f) are displayed. Copy number profile plots show bins (grey dots) and segment (orange) log_2_ copy number ratio (y-axis). The phylogenetic tree leaves (single dissected region) are colored according to histological type and the branches (hamming distances based on CNA segments) annotated with corresponding specific somatic alterations or their total number (CNA: regular, genes: italic font). The tree root corresponds to an inferred normal diploid ancestor. PAM50 subtype of the region is indicated when available. Annotations and trees are available for 10 additional samples in Figure S4.

Substantial heterogeneity and evolutionary patterns are evident in samples like MCL76_061_16200 (Figure 3a-c), where a region of benign columnar alterations preceded two cribriform regions. While all regions shared a WGD event as well as several arm-level CNA and pathogenic mutations in *GATA3* and *SF3B1,* the cribriform region A acquired private 5q and 8q gains and necrotic features. While this example shows tandem genetic and histological changes as seen across the cohort, it also illustrates that despite occurring earlier, the benign region shares many “driver-like” alterations with both cribriform regions. Furthermore, in another example, despite homogeneous cribriform histologies in regions of MCL76_077_15300 (Figure 3d-f) only one cribriform region lost a copy number of chromosome 8, and presented with Her2 PAM50 subtype as opposed to its Luminal A predecessor. Notably bulk studies have shown chromosome 8 loss to be more frequent in Her2 vs Luminal A breast cancers ^35^. Taken all together we illustrate abundant genetic heterogeneity in pure DCIS of all histologies and grades that parallels the levels of phenotypic heterogeneity and often accompanies it, even in regions that are millimeters apart.

### Regional differences in the immune micro-environment

To measure the diversity of the immune-landscape and to investigate its potential association with molecular or histological features, we used multiplex immuno-histochemistry (mIHC) to measure the number and density of four cell types - T-cells (CD3+), B-cells (CD20+), T-regs (CD3+/FOXP3+) and epithelial cells (PanCK+) - according to their proliferative status (Ki67+). Both epithelial (PanCK+) and adjacent stromal (PanCK-proximal to epithelium) areas from pre-malignant (N=36 regions across 32 samples) or benign and normal (N=21 across 21 samples) histologies were evaluated. Among pre-malignant regions, the high-grade epithelial areas had lower cell density due to larger cell sizes and frequent central necrosis (median 3.8 vs 6.4 10^3^ cells/mm2 p<0.03 – Mann-Whitney). Solid lesions had the highest fraction of proliferating epithelial cells (median 11.5% vs 2.8% p<0.02 - Mann-Whitney, Figure 4a), and interestingly 3/10 HG-DCIS cribriform lesions (2 Her2-like, 1 Luminal B) had markedly higher proliferation. We next classified all regions using non-negative matrix factorization of the stromal and epithelial cell densities, resulting in 3 immune-states characterized by their dominant meta-markers (Figure 4b, Figure S5, Table S7): “Active” (ubiquitous high T-cells), “Suppressed” (ubiquitous low T-cells, high B-cells and T-regs) and “Excluded” (high stromal, low epithelial densities). We verified that regions in Excluded state had higher relative stromal cell density, regions in Active state had denser epithelial T-cells and regions in Suppressed state had denser epithelial T-regs and B-cells (Figure 4c-d). A larger fraction of the benign regions were found in Active (7/21) or Suppressed state (9/21) rather than in Excluded state (4/21) and pre-malignant regions in Excluded state were more likely high grade (7/15 vs 2/17 p=0.049). Interestingly, the immune states of benign and pre-malignant regions were concordant in 12/19 matched cases and discordant in 7 whose lesions were specifically in the Excluded state (Figure 4d). This suggests that the Excluded state may be acquired in response to pre-malignant growth, while other states may be intrinsic to various breast micro-environments. Furthermore, pre-malignant regions in Suppressed state were more likely identified in cases younger than 55 (5/8 vs 4/24 OR=7.6 p=0.02), consistent with the younger age of DCIS patients with infiltrating PD-L1+ lymphocytes ^36^. We did not observe any associations between immune states and intrinsic subtype, ER or Her2 status, tumor size, breast density, adipose fraction or DCIS architecture suggesting that they may be independent from traditional histopathological progression risk factors.

**Figure 4.**
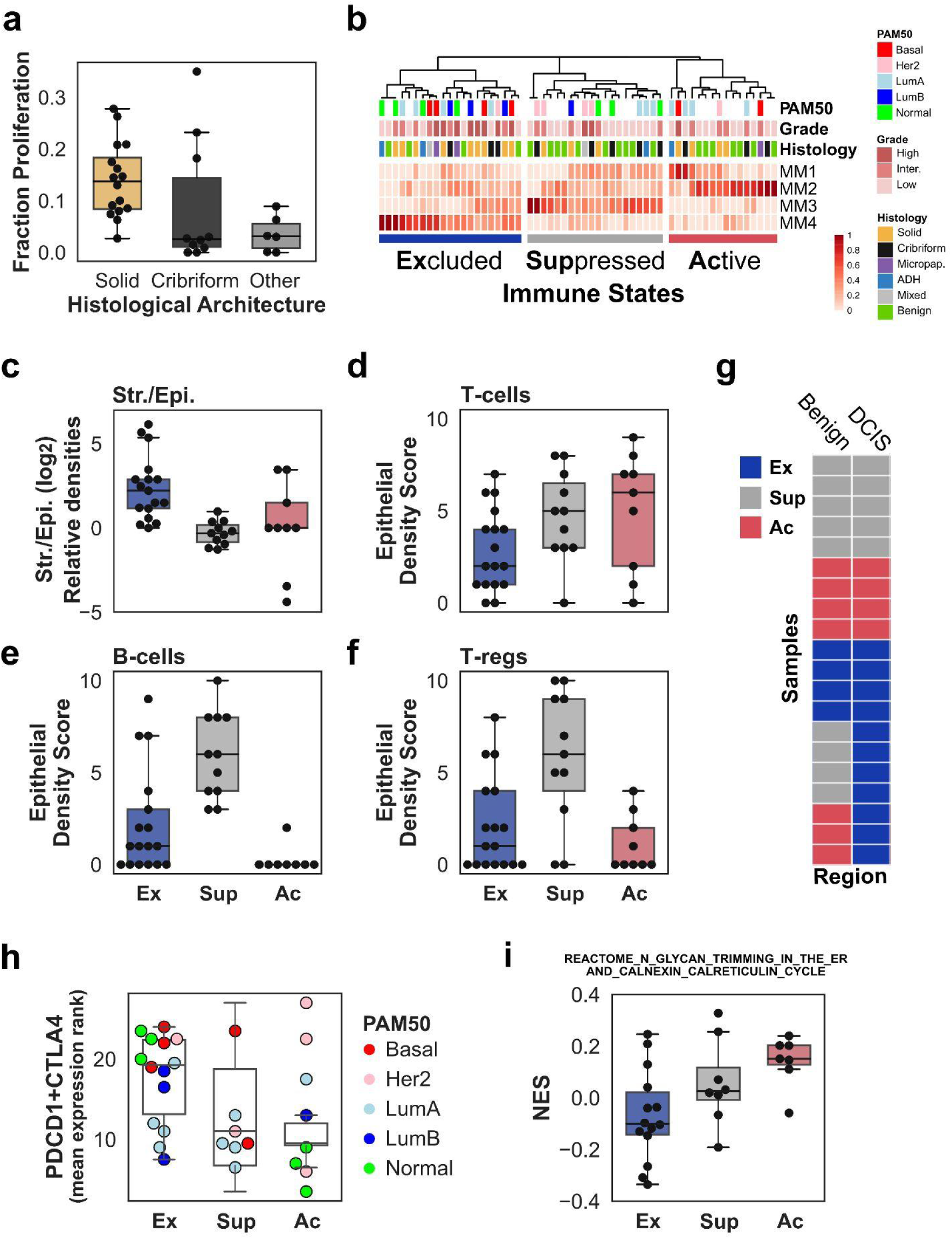
Characterization of the immune landscape. **(a)** Estimates of the fraction of Ki67+ cytokeratin positive cells in epithelium of the mIHC images according to each histological architecture. **(b)** Classification of benign and premalignant regions into 3 immune states according to Meta-Marker (MM) values derived from a non-negative matrix factorization (described in Figure S5). Each state is named after Meta-Marker composition: Active (Ac: MM1&MM2 high TC densities), Suppressed (Sup: MM3 high BC and TREG densities) or Excluded (Ex: MM4 high - low epithelial densities). **(c-f)** Distribution of relative stromal total immune-cell density (c), epithelial cell density scores for T-cells (d), B-cells (e), or T-reg (f) between the 3 immune states. Density scores are obtained from median decile of the cell densities. **(g)** Immune-state comparison in 20 samples (rows) with matching benign (left column) and premalignant (right column) regions. **(h)** Expression of immune checkpoint receptors genes, *PDCD1* and *CTLA-4* in each immune state. **(i)** GSEA normalized enrichment score (NES) for a Reactome gene set across immune states.

In order to identify functional differences between immune-states, we evaluated the differential activity of Hallmark and Reactome processes among the 29 DCIS regions with available gene expression information (Figure S6). Compared to Active and Suppressed states, the Excluded state was associated with upregulation of Type 1 and 2 Interferon response, PD1 signaling and proliferation-related processes as well as the repression of Calcineurin-Calnexin cycle (Figure S6). Noting that the epithelium of DCIS in Excluded state were not completely depleted of infiltrating lymphocytes, the upregulated processes were consistent with the higher expression of *PD1* or *CTLA4* genes in DCIS in Excluded state (Figure 4h), albeit not significant, and suggesting a likely continuum of increasing immuno-suppression from Suppressed to Excluded states. More interestingly, the repression of the Calreticulin-Calnexin cycle was confirmed via single-sample enrichment analysis and showed a progressive repression from Active, to Suppressed, to Excluded states (p=0.022, ANOVA, Figure 4i). This suggests that the export of glyco-proteins - including components of MHC1 complex - via the endoplasmic reticulum, impacts immune-surveillance.

## Discussion

There is a compelling requirement for a DCIS atlas that delivers a relatively unbiased, multi-modal perspective of pre-invasive breast cancer. Here, we report the multi-modal profiling of a diverse set of pure DCIS. This comprehensive atlas both confirms previous molecular findings and provides a higher resolution histological and spatial context to interpret them. However, with only 3 known recurrences, the significance of our observations for progression prognosis could not be formally established. Our findings provide a landscape of representative pure DCIS identified in absence of invasive lesions. While some lesions were small, others were quite extended (N=14 larger than 4 cm), which should capture factors that may be associated with robust containment. The cohort therefore spans a variety of clinical, histological, phenotypic and genotypic features. Such variety and contrast is critical to ensure this atlas’ utility in designing larger studies, or perhaps providing more cautionary interpretation of observations from cohorts enriched for specific risk factors.

At the heart of our study’s innovation was the ability to generate molecular profiles from limited amounts of dissected archival tissue specimen. Similar approaches are used to study clonal expansion in normal tissues ^25, 26^, but generally not performed in parallel for RNA and DNA. Importantly some limitation remains and not all assays were successful. The large variability in success rate was not easy to predict. Likely the age of the specimen, its size, fixation conditions and storage conditions all contribute to success variability which cannot be controlled in a retrospective investigation. Additional limitations are analytical, such as the absence of a matched source of normal DNA from every sample which can result in residual germline variants, perhaps inflating the overall mutation rate observed. The use of adjacent normal or benign tissue can also be problematic and there is ample evidence that they also accumulate somatic mutations ^37^. In our study, we clearly identified known breast cancer driver mutations in samples from ADH or other benign alterations. Overall, while some samples are unlikely to ever contain sufficient material for profiling or dissection of adjacent normal, as methodologies evolve and advance, the success rate and data quality will improve to make molecular premalignant profiling more accessible and as routine as is the case in invasive cancer.

Our report contributes to two major advances for understanding pre-malignant lesions. First, we characterized most samples across four important modalities all within a maximum of 50 µm sequentially sectioned tissue. Such advances were enabled by pre-analytical improvements allowing us to reduce the tissue requirement, to include small lesions, and to precisely match regions of interest across each modality: histology, epithelial gene expression, DNA mutations, and immune landscape. As a result we could isolate regions with different histological features that may coexist within a specimen and more confidently establish their association with expression subtypes, clonal heterogeneity or immune state. For example, the integration of histology and expression subtypes showed clear correlation between cribriform architecture and Luminal A subtype. By integrating histology, expression subtype and immune state we showed that some immune-states are found in benign areas and that there is no clear association between immune state and expression subtype. Hence, the depth and interpretability of the analysis is considerably increased by integrating all modalities at the regional level. This has been clearly the case in large cancer studies such as the TCGA, or, more recently through the integrated analysis of histological and somatic features in normal, aging tissues ^25, 26, 38^. While most studies do not typically include immuno-histochemical or other multiplexed spatial analysis, other important advancements in this field in the past year include spatial proteomics used to evaluate the structure of the myoepithelium in DCIS, and spatial transcriptomics used to identify the transcriptional effect of driver mutations in DCIS, representing the emerging frontier of premalignant tissue characterization ^10, 39^. It is therefore likely that additional spatial profiling compatible with FFPE specimens will bring additional prognostic and mechanistic insights in future DCIS studies.

The other important contribution of our study is the sub-histological analysis to compare regions of interest from the same sample and infer phylogenetic relationships between them. While we determined that the majority of the DCIS samples were classified as Normal-like and Luminal A subtype, typically considered less-aggressive subtypes in breast cancer and reflective of the known precursor stage that DCIS represents, we showed evidence for intrinsic heterogeneity in the PAM50 probabilities, either from the distribution of probabilities within a region or from physically separated regions. This is not entirely surprising as bulk expression subtypes are the result of averaging heterogeneity, similar to glioblastoma subtypes ^40^ or IBC subtypes ^41^ from single-cell analysis. Such heterogeneity, especially in DCIS, had been proposed before on the basis of marker staining ^42^ and our results confirm that it may be rather common. Similarly to the frequency of heterogeneity between region subtypes, we identified evidence of genetic heterogeneity in 7/12 cases, including the presence of private putative driver mutations. This fraction may be an underestimate given the close proximity of many selected pairs. However, the majority of putative genetic drivers, copy number hallmarks and even WGD were clonal, shared by all regions investigated, including a few benign regions. This observation supports evolutionary models derived from invasive cancer, including multi-sample studies, that suggest that most driver mutations occur early followed by a phase of clonal expansion. Similar observations were also made in early multi-regional studies in DCIS ^42–44^ and studies comparing synchronous DCIS-IBC cases using single-cell sequencing ^13^, providing further evidence that breast cancer genetic evolution starts in the pre-invasive stage and possibly in benign regions. It is likely that driver alterations may even be present in adjacent histologically normal tissue as observed in field effects studies in normal tissue and benign ducts ^37, 45^. Such effects support an important contribution of host factors to the initial genetic injury. Hence, unlike previous attempts which were focused on histopathological features, including grade, surgical margins ^46, 47^, future DCIS prognostic models will likely need to be derived from lifetime cancer risk models like GAIL^48^ or BOADICEA ^49^ and incorporate host specific factors, such as polygenic risk scores and reproductive factors, that likely contribute to the DCIS initiation and trajectory.

The immune micro-environment of DCIS has been previously investigated, using both quantification of tumor infiltrating lymphocytes (TIL) and more specific immuno-histochemical approaches and revealed clear quantitative and qualitative variation in lymphocyte infiltration, including higher TIL number and more immunosuppressive features in high risk lesions ^50^. Importantly, previous studies in pure DCIS did not quantify stromal and epithelial TILs separately ^12, 50^. This distinction may be hard to make in IBC, where both compartments interact at the invasive front and pathologist subjectivity can have a major impact ^51, 52^. However, this separation can be more clearly established in the analysis of DCIS and was critical in the identification of the Excluded immune state in our atlas. While the Active and Suppressed states have been observed before and could readily be identified in our data, the identification of the Excluded state required the use of an analytical method (NMF) to account for the strong correlations that can exist between TILs type and compartments. The inclusion of adjacent benign areas was also important to interpret the significance of the immune-states, as the Excluded state appeared more likely in reaction to the DCIS growth and increased grade. The Excluded state exhibited features of immune evasion and could represent a more advanced level of immuno-suppression than the Suppressed state, with the consequence of a topological exclusion from the duct. The downregulation of components of the Calreticulin-Calnexin cycle in the epithelium in Excluded state could impact MHC-I export or maturation providing an evasion mechanism, and contrasting with evasion mediated by MHC-I genetic loss observed in IBC ^53, 54^. It would be interesting to determine whether the immune states identified can explain the variability of response to local injection of anti-PD1 antibody in DCIS patients, and whether any of the states would elicit, or prevent, the desired ductal infiltration by T-cells ^55^.

As illustrated by our study and recent advances in the profiling of normal tissues ^25, 26^, histopathology and molecular pathology are becoming more integrated fields, generating deeper and broader datasets at increased cellular and spatial resolution, from the most challenging human samples. Future studies of early transformation and pre-cancer biology such as the one presented here will likely benefit the most from such approaches which capture heterogeneity at scale and can help reconcile analog (optical) and digital (genomics and multiplex) observations. As a result, such multi-dimensional integration may help identify common factors mediating epithelial transformation and progression across multiple glands and organs.

## Methods

### Sample collection and preparation

FFPE blocks were obtained from UCSD or UVM Pathology Departments after surgical biopsy, excision or mastectomy. The study was reviewed and approved by each institutional review board and they granted a waiver of consent. All specimen blocks were de-identified and sectioned sequentially for the following purpose: Hematoxylin-Eosin (H&E) staining (N=1; 4 µM glass slide), Laser Capture Microdissection (LCM; N=3; 7 µM glass slide coated with polyethylene naphthalate – ThermoFisher #LCM0522), multiplex or regular immunohistochemistry (N≥3 4 µM glass slide) and a final H&E staining (N=1; 4 µM glass slide). The H&E slides were scanned at high resolution and reviewed and annotated by the study pathologist. The LCM slides were stored at −20°C in an airtight container with desiccant until ready for dissection (1 day to 3 months). The LCM sections were thawed and stained with eosin, sections were kept in xylene and dissected within 2 hours of staining. LCM was performed using the ArcturusXM Laser Capture Microdissection System (ThermoFisher). Matching regions from 6 adjacent sections were collected on capsure Macro Cap (for DNA, N=3 slides) or HS caps (for RNA, N=3 slides), region size, and unambiguous match permitting. Post-dissection, all caps were covered and stored at −20°C with desiccant. *<u>DNA extraction and QC:</u>* The membrane and adhering tissue were peeled off the caps using a razor blade and the peeled membrane was incubated in proteinase K digestion reaction overnight for 16 h at 56°C to maximize DNA yield after cell lysis The DNA was extracted using the QIAamp DNA Micro Kit (Qiagen) and the elution was done in 20 µL. The extracted DNA was quantified by fluorometry (HS dsDNA kit Qbit – Thermofisher).

### RNA-Sequencing and analysis

#### Library Preparation

RNA sequencing was performed using SMART-3Seq, a 3’ tagging strategy specifically designed for degraded RNA directly from FFPE LCM specimen ^24^. LCM dissected SMART-3Seq libraries were prepared using the standard protocol for FFPE tissue on Arcturus HS LCM Cap and the individual library SPRI purification option. All FFPE LCM dissected libraries were amplified using 19 PCR cycles during indexing to minimize over-amplification of high abundance mRNAs in each library. Libraries were individually analyzed for size distribution on an Agilent 2200 TapeStation with High Sensitivity D1000 reagent kits to verify average library size of 190 bp and stored at −20 C until sequencing. When all libraries were ready for sequencing, 1 µL of each library was then used to create two library pools used for sequencing and quantified by Qubit 2.0 Fluorometer HS DNA assay. Library pools were sequenced with a 1% PhiX spike-in control library and sequenced on an Illumina HiSeq4000, a run type of single read 75 (SR75) and dual index sequencing.

#### Transcriptome analysis

Read count data was obtained using a dedicated analysis workflow https://github.com/danielanach/SMART-3SEQ-smk. Briefly, sequencing reads were trimmed using cutadapt 1.18, UMIs were processed using the umi_homopolymer.py script in the SMART-3SEQ tools (https://github.com/jwfoley/3SEQtools), aligned using STAR 2.6.1a, deduplicated using the dedup.py script from https://github.com/jwfoley/umi-dedup and read counts were calculated using featureCounts 1.6.3 ^56, 57^. Count data was then merged and filtered to remove samples with fewer than 55,000 counts and genes with fewer than 10 read counts across all samples. Filtered count data was then loaded into Seurat version 3.2.3 and processed using the *SCTransform()* function version 0.3.2 to regress out the high mitochondrial content variability across the samples ^58^. Batch correction was then performed using ComBat to remove variation attributable to the sequencing center (UCSD vs UVM) ^59^. PAM50 subtype probabilities were calculated from the *SCTransform* and batch normalized data using the genefu package ^60^. Gene set enrichment analysis (GSEA) was performed as in ^61^ and single-sample GSEA as in ^62^. Gene sets from the REACTOME and Hallmark collections in MSigDB were used to compare the excluded to the non-excluded groups, a permutation test was performed to assess the significance of the GSEA results ^63, 64^. ANOVA was used to compare the ssGSEA results between the three mIHC groups. FDR of less than 0.1 and p-values of less than 0.05 were considered significant.

### Whole exome sequencing and primary analysis

#### LIbrary preparation

DNA was sheared down to 200 base pairs (bp) using Adaptive Focused Acoustics on the Covaris E220 (Covaris Inc) following manufacturer recommendations with 10 μL Low EDTA TE buffer supplemented with 5 μL of truSHEAR buffer using a microTUBE-15. Libraries were prepared using the Accel-NGS 2S PCR-Free DNA Library Kit (Swift Biosciences). Ligated and purified libraries were amplified using KAPA HiFi HotStart Real-time PCR 2X Master Mix (KAPA Biosystems). Samples were amplified with 5 μL of KAPA P5 and KAPA P7 primers. The reactions were denatured for 45 seconds (sec) at 98°C and amplified 13-15 cycles for 15 sec at 98°C, for 30 sec at 65°C, and for 30 sec at 72°C, followed by final extension for 1 min at 72°C. Samples were amplified until they reached Fluorescent Standard 3, cycles being dependent on input DNA quantity and quality. PCR reactions were then purified using 1x AMPure XP bead clean-up and eluted into 20 μL of nuclease-free water. The resulting libraries were analyzed using the Agilent 4200 Tapestation (D1000 ScreenTape) and quantified by fluorescence (Qubit dsDNA HS assay).

#### Capture and Sequencing

Samples were paired and combined (12 μL total) to yield a capture “pond” of at least 350 ng, and supplemented with 5 μL of SureSelect XTHS and XT Low Input Blocker Mix. The hybridization and capture was performed using the Human All Exon V7 panel (S31285117) paired with the Agilent SureSelect XT HS Target Enrichment Kit following manufacturer’s recommendations. Post-capture amplification was performed on the beads in a 25 μL reaction: 12.5 μL of nuclease-free water, 10 μL 5x Herculase II Reaction Buffer, 1 μL Herculase II Fusion DNA Polymerase, 0.5 μL 100 millimolar (mM) dNTP Mix and 1 μL SureSelect Post-Capture Primer Mix. The reaction was denatured for 30 sec at 98°C, then amplified for 12 cycles of 98°C for 30 sec, 60°C for 30 sec and 72°C for 1 min, followed by an extension at 72°C for 5 minutes and a final hold at 4°C. Libraries were purified with a 1x AMPure XP bead clean up and eluted into 20 μL nuclease free water in preparation for sequencing. The resulting libraries were analyzed using the Agilent 4200 Tapestation (D1000 ScreenTape) and quantified by fluorescence (Qbit – ThermoFisher). All libraries were sequenced using the HiSeq 4000 sequencer (Illumina) for 100 cycles in Paired-End mode. Libraries with distinct indexes were pooled in equimolar amounts. The sequencing and capture pools were later deconvoluted using program bcl2fastq [19].

#### Sequencing reads processing and coverage quality control

Sequencing data was analyzed using bcbio-nextgen (v1.1.6) as a workflow manager [20]. Samples prepared with identical targeted panels were down-sampled to have equal number of reads using seqtk sample (v1.3) [21]. Adapter sequences were trimmed using Atropos (v1.1.22), the trimmed reads were subsequently aligned with bwa-mem (v0.7.17) to reference genome hg19, then PCR duplicates were removed using biobambam2 (v2.0.87) ^65–67^. Additional BAM file manipulation and collection of QC metrics was performed with picard (v2.20.4) and samtools (v1.9) ^68^. The summary statistics of the sequencing and coverage results are presented in Table S8.

### Identification of somatic mutation and copy number alterations

#### Variant calling

Single nucleotide variants (SNVs) and short insertions and deletions (indels) were called with VarDictJava (v1.6.0), and Mutect2 (v2.2) ^69, 70^. Variants were required to fall within a 10 bp boundary of targeted regions that overlapped with RefSeq genes (v 109.20190905). A pool of normal DNA was created using whole exome sequencing data of blood of 18 unrelated individuals and was used to eliminate artifacts and common germline variants. Only variants called by both algorithms were considered. These variants were then subjected to an initial filtering step with default bcbio-nextgen tumor-only variant calling filters and the following parameters were used: position covered by at least 5 reads, mapping quality more than 45, mean position in read greater than 15, number of average read mis-matches less than 2.5, microsatellite length less than 5, tumor log odds threshold more than 10, Fisher strand bias Phred-scaled probability less than 10 and VAF more than 0.1 ^71^. Functional effects were predicted using SnpEff (v4.3.1) ^72^. All samples were re-evaluated for the presence of COSMIC (v91) database mutations which have been previously observed in at least 15 patients and fall within 137 known breast cancer driver genes (Table S9) ^7^.

#### Germline variant filtering

In absence of matched normal tissue for DCIS samples, somatic mutations were prioritized computationally using the approach from the bcbio-nextgen tumor-only configuration then additionally subjected to more stringent filtering ^71^. Briefly, common variants (MAF>10^-3^ or more than 9 individuals) present in population databases - 1000 genomes (v2.8), ExAC (v0.3), or gnomAD exome (v2.1) - were removed unless in a tier 1 gene from the cancer gene consensus and present in either COSMIC (v91) or clinvar (20190513) ^7, 73–76^. Variants were removed as likely germline if found at a variant allelic fraction (VAF) greater or equal to 0.9 in non-LOH genomic segments – as determined by CNA analysis (below). Lastly, variants were also removed as potential germline (or artifact) if found in more than two patients in the pool of normal (described above).

#### Single-sample CNA calling

CNVkit ^77^ was used for calling somatic copy number alterations (CNA) to measure both overall CNA burden, arm and gene level CNA and identify LOH as previously described in ^23^. Allele-specific copy number calling algorithm, ASCAT, was used on a select number of samples for which there was sufficient coverage and the algorithm converged on a solution, in order to identify whole-genome doubling events as well as confirm CNA identified by CNVkit ^78^. Default parameters were used with ASCAT with the exception of a segmentation penalty of 100 and a gamma of 1.

#### Multi-region CNA segmentation

To generate harmonized segmentation breakpoints between regions belonging to the same sample, multi-region segmentation as performed with the R CopyNumber (v1.26.0) package ^79^. Outliers in CNVkit bin-level log2 copy ratios were detected and modified using Median Absolute Deviation Winsorization with the winsorize() function, segments were then called using the *multipcf()* function with a gamma of 40.

#### Mutational signatures

Mutational signatures were called on merged region samples using a single-sample variation of SigProfiler with default parameters to decompose into known single-base substitutions (SBS) reported in COSMIC ^80, 81^.

### Analysis of the clonal evolution and genetic heterogeneity

#### Measurement of genetic divergence

Divergence was measured on each pair of related regions, *a* and *b*, using the following equation:

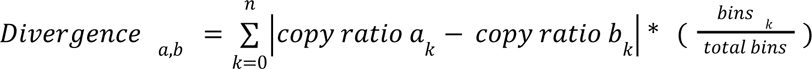

Where *k* is the copy number segment, *n*is the total segments, and *bins*is the number of bins covered by a segment from the CNVkit input file. The 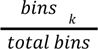 term was used as a weighted correction factor for the number of bins contributing to a segment. For samples with more than 2 regions, the maximum divergence between any two regions was used to represent the sample.

#### CNA-based phylogenetic reconstruction

Construction of phylogenetic trees was performed similarly to the methodology outlined in ^82^. Briefly, for each sample the log2 copy ratios from multi-sample copy number segments with at least 12 probes (see above), were translated into a matrix containing −1 for loss (log2 copy ratio<-0.6), 0 for neutral (−0.4<= log2 copy ratio <=0.3) and undetermined for anything else. This matrix was then used to generate Maximum Parismony trees using phangorn using default parameters ^83^.

#### Mutation-based phylogenetic reconstruction

To allow the analysis of clonal relationships between regions of the same sample, the coverage depth of each allele at any remaining mutated position in any region was extracted using Mutect2 joint variant caller on the sets of aligned reads from each region. In order to call a mutation either absent or present in a region, we used a Bayesian inference model specifically designed for multi-region variant calling ^84^. Treeomics (v1.7.10) was run with the default parameters except for e=0.02. The tree solution which matched the CNA-based reconstruction was then integrated into a single tree for Figures 3 and S4.

### Multiplex Immuno-histochemistry

#### Staining

Tissue sections were prepared from formalin-fixed paraffin embedded tissue blocks and cut to 4 micrometers serial sections and mounted on Superfrost Plus (VWR). The procedure for multiplex immunohistochemistry (mIHC) was followed by a manufacturer’s protocol for Opal7-color automation IHC kit (Akoya Bioscience), and the staining was performed with Autostainer DISCOVERY ULTRA (Ventana). Antibodies used in mIHC are anti-CD3 (clone 2GV6, Ventana), anti-CD20 (clone L26, Ventana), anti-Ki67 (clone 30-9, Ventana), anti-FOXP3 (clone SP97, Spring), anti-pan cytokeratin (CK; clone AE1/AE3, DAKO), anti-CD117 (clone c-kit, DAKO). The molecular markers of immune panel (CD3, CD20, Ki67, CKs, FOXP3 and CD117) were visualized with Opal520, Opal540, Opal570, Opal620, Opal650 and Opal690, respectively. DAPI counterstaining was performed with Discovery QD DAPI (Roche). ProLong Diamond Antifade Mounting (ThermoScientific) was used for mounting the coverslip. Detailed staining conditions and autostainer’s protocols are reported in our recent report ^85^.

#### Visualization and analysis

Tissue samples stained with mIHC were scanned with multispectral imaging microscopy (Vectra 3, Akoya Bioscience). Scanned multispectral images were unmixed on inForm software (ver.2.4.0, Akoya Bioscience) to acquire the fluorescence signal from each marker ^85^. Imaging analysis was performed on inForm software by identifying tumor (CK+ area) and stroma (CK-area proximal to the epithelium), each nucleated cell and its cell type. Alternatively, QuPath software ^86^ was also used to perform similar imaging analysis on unmixed images converted to multi-layered TIFF format by inForm software ^85^. Scanned image areas were aggregated into up to three histological regions per sample: main pre-invasive lesion, alternate pre-invasive lesion, benign or normal epithelium. In each region, the stromal and epithelial densities of each cell type and state was calculated, including when cells were not present (density=0). Regularized marker densities into distribution deciles were then used to classify samples using non-negative matrix factorization (Table S7). The immune-states were assigned and named after the hierarchical clustering of the H matrix (meta-marker values).

### Whole slide image digital analysis

High resolution whole slide images of H&E stains were loaded into a QuPath (v2.3) project ^86^. One analysis area was defined for each specimen, avoiding location of biopsies as well as dust or marked areas. The analysis areas were segmented into superpixels (sigma=5 µm, spacing=50 µm, maxIterations=10, regularization=0.25) and each superpixel was annotated with both Hematoxylin and Eosin Intensity features (size=2 µm, tile size=25 µm). The mean, median, min, max and standard deviation values were then smoothed (Haralick distance=1, Haralick bins=32). Multiple training areas were annotated from each of the following classes: adipose, stroma, inflammation, epithelium (normal and atypical), void, necrosis, blood vessels. Multiple areas across 2 to 4 samples were used to train a Random Tree classifier. The classifier was then applied to all superpixels included in the analysis area. The accuracy of the classifier was assessed both visually and with multiple test areas for each class. Superpixels of the same class were merged into single annotations and the resulting areas recorded. Separate classifiers were used for images from different institutions, to mitigate possible variation staining, scanning or image format. The fraction of adipose area was compared to breast density using Mann-Whitney test comparing dense & heterogeneously dense breast to other lower densities, or comparing solid DCIS to non-solid DCIS lesions.

## Data Availability

The raw RNA and DNA sequencing data has been deposited in dbGAP phs002225. High resolution whole slide images of the H&E stains and corresponding annotations can be viewed on the JPL LabCAS portal (digital object identifiers included in the Table S1).

## Acknowledgments

We are grateful to Drs. Michael Campbell, Christina Yau and Kathleen Curtius for helpful conversations, Drs. Alfredo Molinolo, Oluwole Fadare, Sharmeela Kaushal, Valeria Estrada and Mrs Kimberly Mcintyre for their support and assistance in the tissue collection, preparation and dissection, Mrs Eliza Jeong, Marcy Andersen and Nicole Lee for their assistance retrieving clinical information. We thank the technical assistance of the Vermont Integrative Genomics Resource Massively Parallel Sequencing Facility with the combined support of the University of Vermont Cancer Center, Lake Champlain Cancer Research Organization, UVM College of Agriculture and Life Sciences, and the UVM Larner College of Medicine. We acknowledge the work of all working groups from the NCI Consortium for the Molecular Characterization of Screen Detected Lesions (MCL), in particular Christopher Amos, Daniel Crichton, Heather Kincaid, Kirsten Anton, Luca Cinquini, David Liu for their assistance in the data management and sharing. Figure 1 was created with BioRender.com and printed with permission.

This work is supported by funding from the National Institute of Health (U01CA196406, U01CA196406-03S1, U01CA196383, R01DE026644, T32GM008806, T15LM011271), the National Cancer Institute (P30CA023100) and the California Tobacco Related Disease Research Program pre-doctoral fellowship to DN (28DT-0011). The funding bodies had no role in the design of the study; collection, analysis, and interpretation of data; or in the writing of the manuscript.

## Authors Contributions

JLS, GS, GLH, LJE, SDB, OH designed the study, FH, AM, SDB, DLW, BLS, TOK selected and annotated the specimen. JS, KJ, HZ, HM, MFE, JG generated the data, OH, DN, AO, HM analyzed the data, OH and DN wrote the manuscript. All authors reviewed and approved the manuscript.

## Supplemental Figures

**Figure S1:**
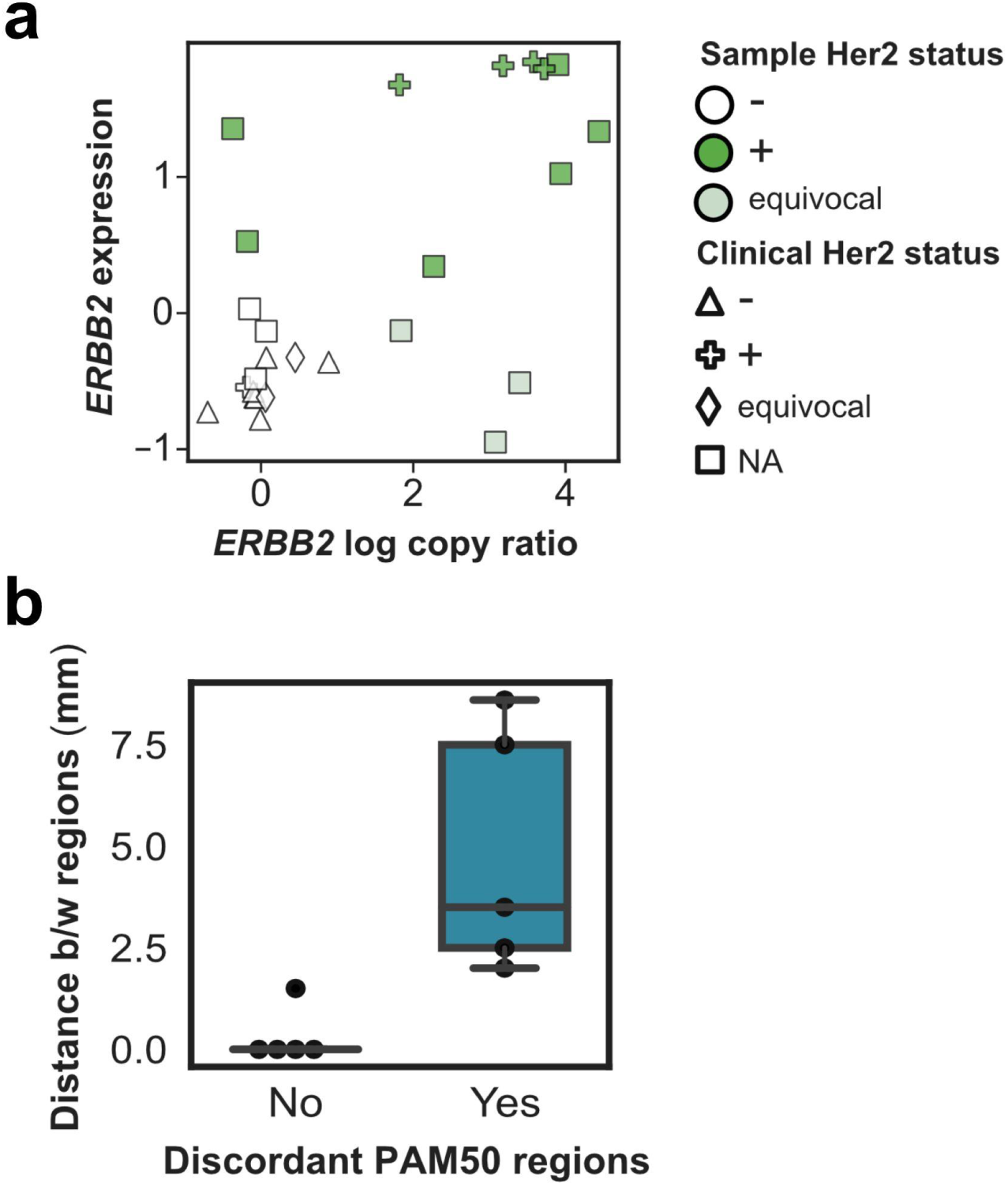
Pure DCIS characterization. **(a)** Estimation of Her2 status. The DNA-based log_2_ copy number ratio (x-axis) and RNA-based expression level (y-axis) of *ERBB2* gene are displayed for 26 samples with both data available. **(b)** PAM50 discordance between regions in relationship with spatial distance between regions.

**Figure S2.**
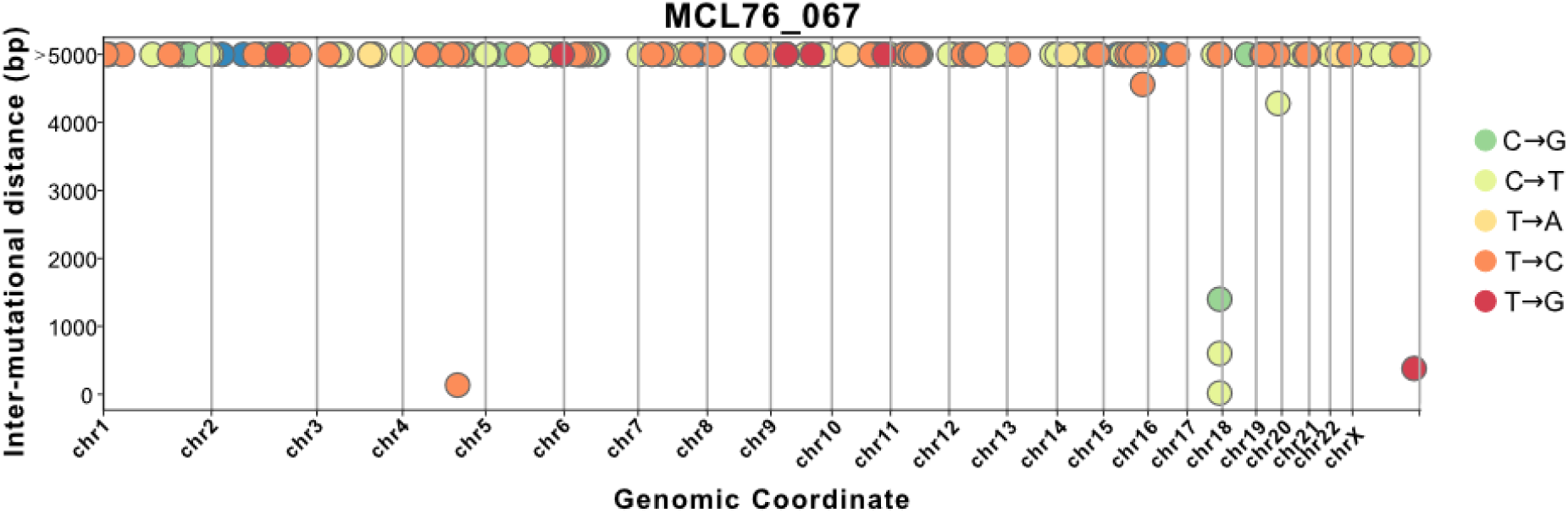
Likely kataegis event in MCL76_067_16600 in chromosome 17. Along the genome coordinate (x-axis), the relative distance between proximal mutations (y-axis) as well as their substitution type (colors) are indicated.

**Figure S3.**
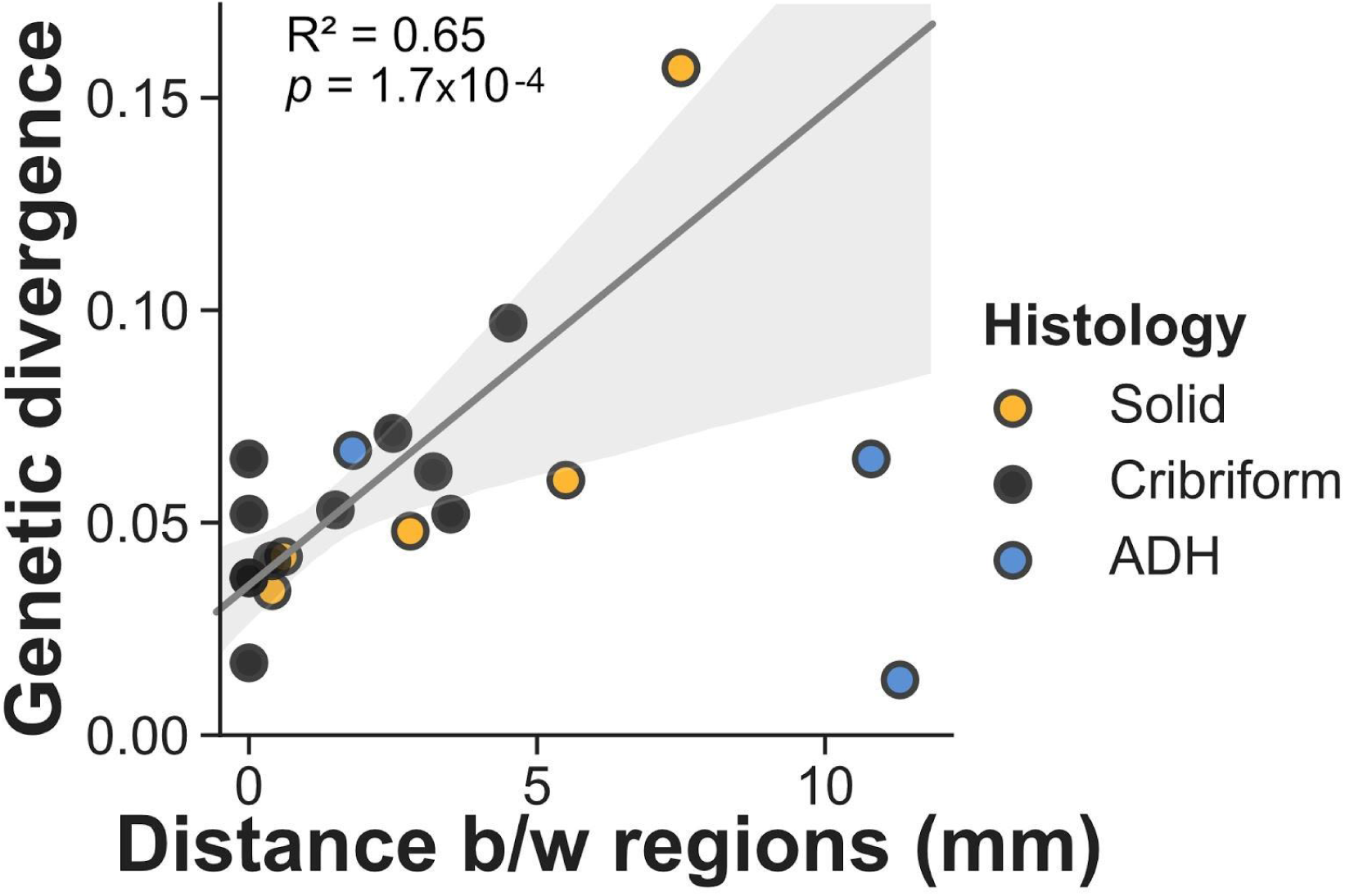
Genetic divergence in multi-region DCIS. CNA-based genetic divergence (y-axis) of each pair of histologically concordant regions (dot) as a function of the minimum physical distance between them (x-axis), colored by their histology. Linear regression line fit of DCIS samples shown in solid dark gray line, with 95% confidence interval estimate based on bootstrapping in light gray.

**Figure S4.**
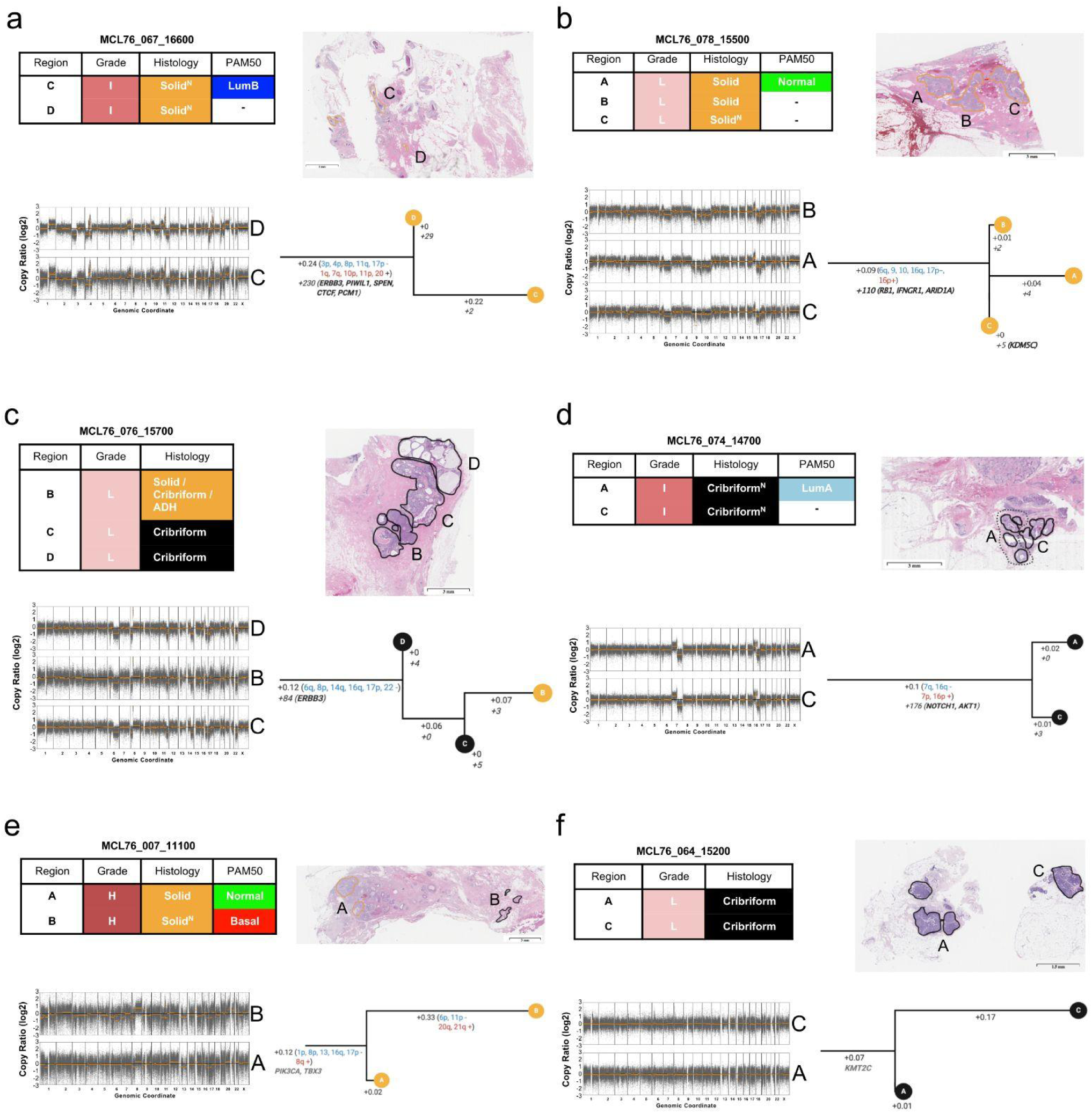
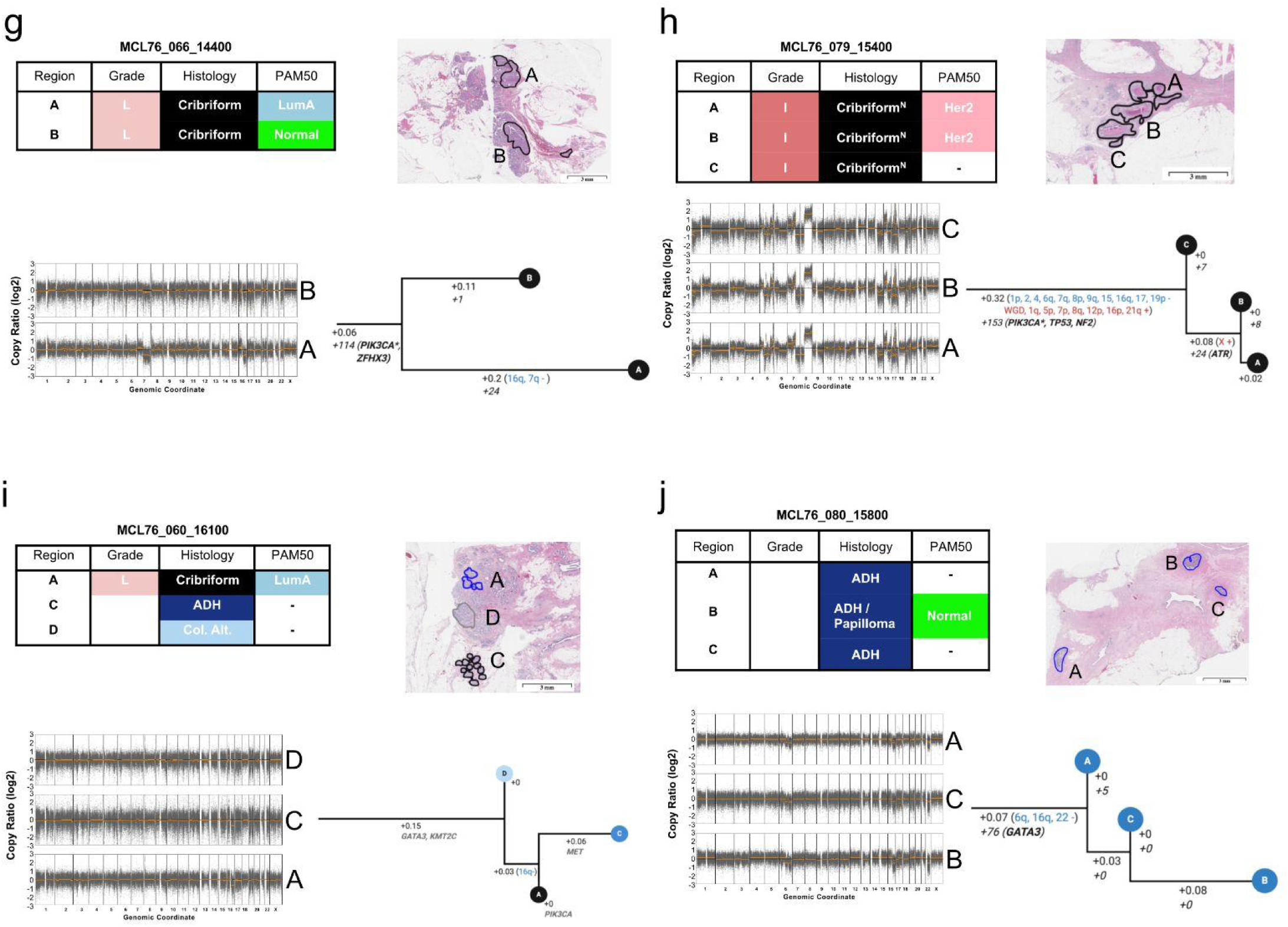
Phylogenetic trees for multi-region DCIS samples. **(a-j)** Clonal reconstruction using CNA and somatic mutations in 25 related regions across 10 samples. In each panel, *<u>top left:</u>* a table describing the name, nuclear grade and histological architecture of each region in a sample is shown, (necrosis is indicated with N); *<u>top right:</u>* shows an H&E image of the sample with dissected regions drawn on the image; *<u>bottom left</u>*: Copy number profiles for each region in the sample, genomic coordinates are indicated on the X-axis and the log_2_ copy ratio on the Y-axis. Bins are indicated in dark-grey and segments in orange; *<u>bottom right</u>*: Phylogenetic tree for the sample with leaf nodes indicating a single dissected region colored by histology. The tree is rooted to a normal diploid ancestor. Branch lengths are hamming distances based on CNA segments. Branches are labeled with 1) CNA-based branch length, and, when available, 2) arm-level CNA losses (blue) and gains (red) and 3) somatic mutation number (italic) with mutations in breast cancer driver genes indicated (black: high coverage, grey: low coverage). CNA smaller than arms, or on driver genes are not displayed.

**Figure S5.**
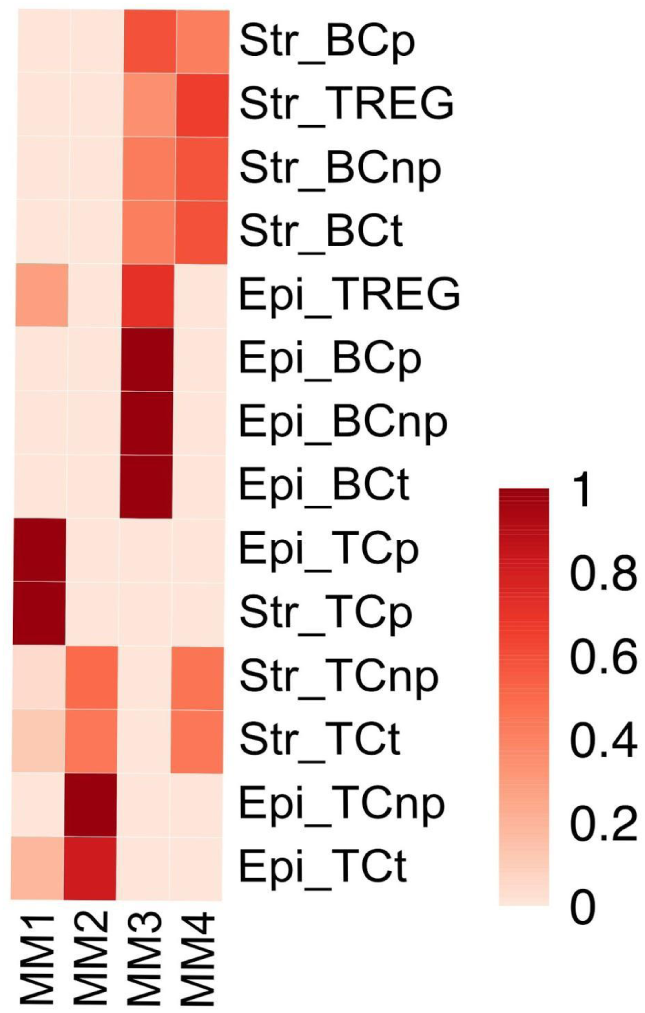
Composition of the NMF Meta-markers (columns MM1-4) according to the densities scores (red scale) of each cell type (BC: B-cells, TC: T-cells, TREG: regulatory T-cells), proliferative state (p: Ki67+, np: KI67-, t: total) and regional location (Epi: Epithelium, Str: Stroma). Estimates of the fraction of Ki67+ cytokeratin positive cells in epithelium of the mIHC images according to each histological architecture.

**Figure S6:**
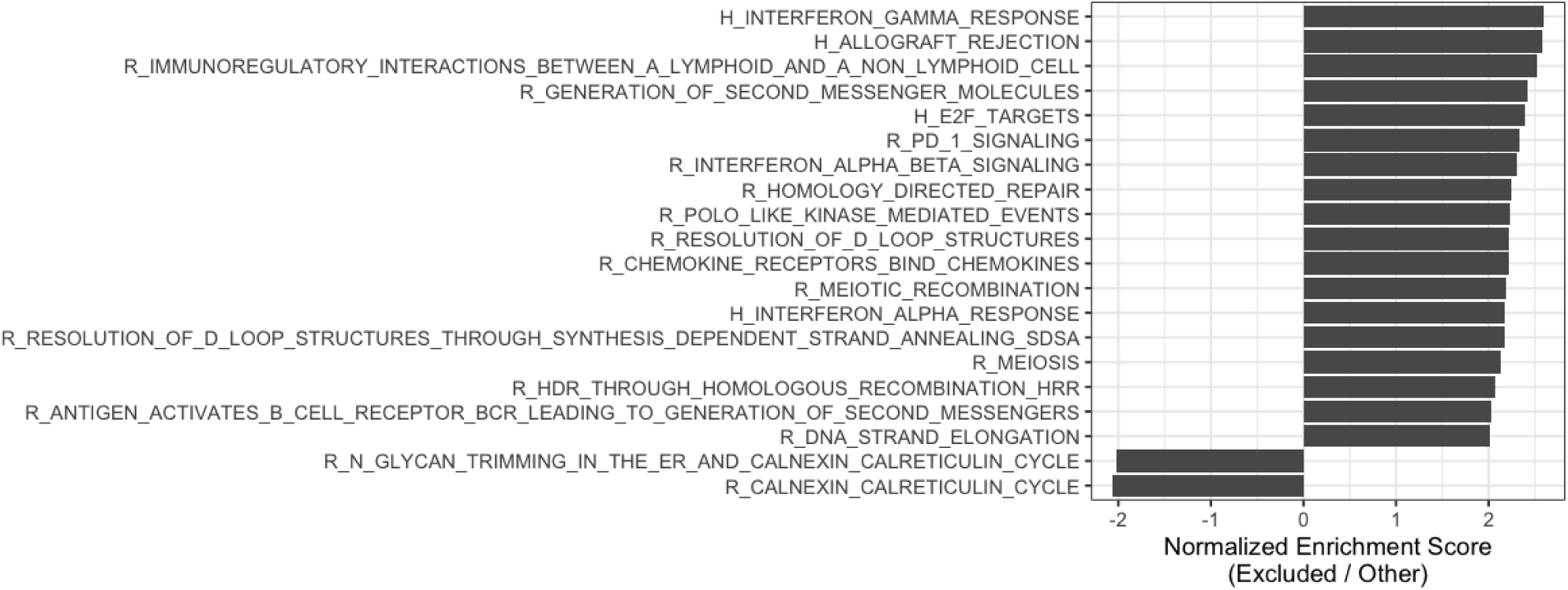
Gene sets significantly deregulated in epithelium of regions in the Excluded immune state. All Hallmark (H) and Reactome (R) genesets were tested. Genesets with an absolute normalized enrichment score greater than 2 and with FDR less than 0.05 are represented.

